# Colonization of a novel host plant reduces phenotypic variation

**DOI:** 10.1101/2023.01.16.524271

**Authors:** Kalle J Nilsson, Masahito Tsuboi, Øystein Opedal, Anna Runemark

## Abstract

Understanding the evolutionary potential of populations –evolvability– is key to predicting their ability to cope with novel environments. Despite growing evidence that evolvability determines the tempo and mode of adaptation, it remains unclear how adaptations to novel environments influence evolvability in turn. Here we address the interplay between adaptation and evolvability in the peacock fly *Tephritis conura*, which recently underwent an adaptive change in the length of female ovipositor following a host shift. By comparing evolvability in various morphological traits including female ovipositor length between ancestral and derived host races, we found that evolvability is decreased in females of the derived host race compared to the ancestral host race. We found a correlation between evolvability and divergence between populations in both sexes, indicating that the overall pattern of evolvability has not been disrupted by the host shift despite the reduction in females of the derived host race. Exploration of the pattern of phenotypic integration further revealed that the ovipositor length constitutes a module that is separated from other measured traits. These results suggest that adaptation to novel environments can affect evolvability, and that modularity helps minimizing detrimental effects that adaptations may cause to other correlated traits.

## Introduction

Variation is the raw material for adaptive evolution. Natural selection results from variation in genotypes, phenotypes and fitness, and the evolutionary potential for response to selection is determined by standing genetic variation. There is mounting evidence that a lack of evolutionary potential may constitute evolutionary constraint to adaptive divergence (Bradshaw and McNeilly 1991; Schluter 2000; Arnold et al. 2001; Bolstad et al. 2014; Houle et al. 2017; McGlothlin et al. 2018), and that evolutionary potential is highly variable across traits and species (Hansen and Pelabon 2021). A key question is therefore how evolutionary potential evolves, and how this interacts with patterns of adaptation during the process of evolutionary divergence (Berner et al. 2010; Eroukhmanoff and Svensson 2011).

Quantitative genetics provides a powerful approach to quantifying the evolutionary potential of a species. Phenotypic variation plays a prominent role in the theoretical framework of evolutionary quantitative genetics, as summarized by the simple ‘Lande equation’ (Lande 1979). This framework emphasizes the role of the additive genetic variance-covariance matrix (**G**) as a key determinant of response to selection, i.e. evolutionary potential. If the G-matrix remains relatively stable over time, adaptive evolution could be well understood through focusing on the dynamics of selection. However, if **G** itself evolves (Jones et al. 2003; Arnold et al. 2008; Milocco and Salazar-Ciudad 2022) and changes over time, the predictive power associated with a contemporary estimate of **G** is critically dependent on our understanding of the dynamics of **G** (Walsh and Blows 2009; McGlothlin et al. 2018).

Previous work suggests that **G** does evolve in natural populations (Cano et al. 2004; Doroszuk et al. 2008; Björklund et al. 2013; Walter et al. 2018), but how **G** evolves is less well understood. This is partly due to the difficulty in formulating testable hypotheses regarding the evolution of **G** (Pélabon et al. 2010). For example, the structure of variance-covariance matrices may be altered following selection (Revell et al. 2010; Penna et al. 2017) and ancestral bottlenecks are expected to affect current evolvability by reducing genetic variation (Nei et al. 1975). However, a bottleneck may also increase genetic variance if there is cryptic genetic variance under effects of epistasis, dominance or the environment (Whitlock et al. 2002; Paaby and Rockman 2014). Gene flow among diverging lineages can also increase evolvability if mixed genetic variants result in phenotypic variation that is relevant for selection (Guillaume and Whitlock 2007).

The examples above illustrate the complexity of deriving general predictions based solely on first principles. To narrow down the parameter space, empirical studies are needed that compare variational properties of diverging populations with known histories of selection. Host shifts provide an excellent opportunity in this regard because we know *a priori* that ancestral and derived populations are evolving towards different phenotypic optima (Assis et al. 2016). Here, we take advantage of a recent host shift in the peacock fly *Tephritis conura* (Diegisser et al. 2006a; Diegisser et al. 2006b, 2007; Diegisser et al. 2008; Nilsson et al. 2022; Ortega et al. *[in prep*.]) to study evolution of variation during population divergence.

Adult *T. conura* tephritid flies oviposit into the buds of *Circium* thistle buds, and larva and pupae develop within the buds. The ancestral host plant is *Cirsium heterophyllum*, but a subset of populations has recently undergone a host shift from *C. heterophyllum* to *C. oleraceum* (Romstock-Volkl 1997). Interestingly, there is evidence of phenotypic adaptation to the specific host plants, most clearly in the length of the ovipositors. Flies infesting *C. oleraceum* have shorter ovipositors than flies infesting *C. heterophyllum* (Diegisser et al. 2007; Nilsson et al. 2022), matching the smaller bud size of the plant (Romstock-Volkl 1997). Moreover, there is empirical evidence for strongly reduced survival on the alternative host plant (Diegisser et al. 2008), suggesting strong host plant-mediated selection.

The documented natural history of our *T. conura* populations allows us to empirically examine the evolution of variation over the course of population divergence. We approach this question from three perspectives. First, for traits that differ between the ancestral and derived host race, we expect historical and potentially current directional selection in the derived host race. Specifically, selection on the ovipositor to match the bud size of the derived host plant (Romstock-Volkl 1997) likely caused directional selection in the derived host race which has a shorter ovipositor length compared to the ancestral host race (Diegisser et al. 2007; Nilsson et al. 2022). This selection may have altered the structure of the phenotypic variance covariance matrix (**P**) in the derived host race in *T. conura*. Using the concept of conditional variance and autonomy that quantifies the degree of independence among correlated traits (Hansen et al. 2003), we evaluate how the hypothesized directional selection on the ovipositor in the derived host race of *T. conura* affects the overall structure of the P-matrix.

Second, we explore if the evolvability in populations that exist in sympatry with the alternative host race is higher than in allopatric populations due to gene flow between the host races. Secondary sympatry between two recently diverged conspecifics has been suggested to impact the phenotypic covariance as gene flow could increase the combinations of traits available to selection (Blows and Higgie 2003; Dochtermann and Matocq 2016). Host races specializing on the two different host plants coexist geographically in a broad zone where the southern *C. oleraceum* and the northern, ancestral, *C. heterophyllum* are both common (Fig. 1A). This enables us to test to which extent coexistence with the other host race affects **P**.

**Figure 1.**
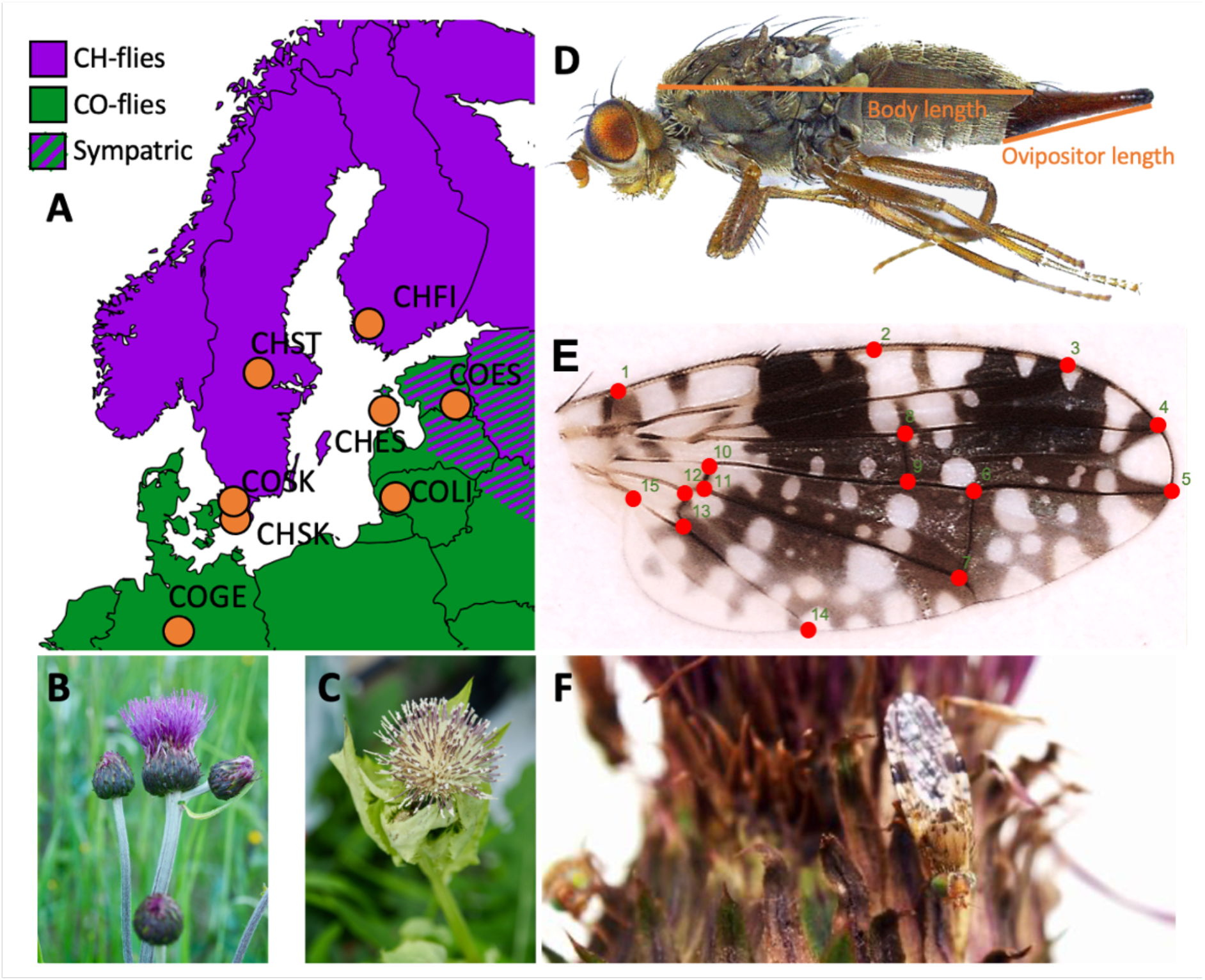
Sampling design, host plants, and traits investigated. **A)** Parallel sampling of allopatric and sympatric populations of the two host races of *T. conura* flies east and west of the Baltic. **B)** The ancestral host plant, *C. heterophyllum*. **C)** The derived host plant, *C. oleraceum*. **D)** Morphometric size measurements of *T. conura*. **E)** Landmarks used for wing shape morphometrics of *T. conura*. **F)** *Tephritis conura* on a *C. heterophyllum* bud.

Finally, we evaluate whether patterns of divergence are themselves correlated with the structure of the ancestral P-matrix, thus testing the hypothesis that ancestral variation can constitute genetic constraints (Bolstad et al. 2014; Houle et al. 2017; McGlothlin et al. 2018). We assess this by asking whether patterns of host-race divergence align with variation within the ancestral host race, so that the host races are diverging in a direction of greater-than-average ancestral variation (Schluter 1996).

## Methods

### Sampling

To examine the distribution of phenotypic variation within and among fly populations specialized on the derived and ancestral host plant we sampled flies from four populations of each host race (Fig. 1A). A haplotype analysis suggests that the host shift took place during the last ice age (∼18 thousand years ago) in the Alps (Diegisser et al. 2006b). While alternative host races are largely reproductively isolated due to differences in the location at which copulation takes place and differences in phenology (Romstock-Volkl 1997), there is evidence of gene flow between the two host races (Diegisser et al. 2006b; Ortega et al. *[in prep*.]). To allow assessing a potential effect of gene flow on evolvability, we included allopatric populations as well as populations that were regionally sympatric with the other host race. We collected thistle buds infested by *T. conura* during the pupal stage at all these locations (Fig. S1) and allowed adults to eclose in a common laboratory environment as described in Nilsson et al. (2022). The sampling scheme enables us to examine to what extent patterns of phenotypic variance are explained by host plant adaptations and by regional sympatry with the other host race. We use the terms sympatric and allopatric to refer to the presence of one or both thistle species on a regional scale (Fig. 1A).

### Measures of morphology and wing shape

After collecting *T. conura* adults, one female and one male per bud was euthanized by freezing individuals for a few days after eclosion, and subsequently included in the morphological analysis. We chose ovipositor length, several size measurements, wing melanisation, and a large set of wing shape measurements as study traits. The ovipositor length is a key functional trait, and we thus consider it separately form the size and shape measurements. Previous work with fly wings of Drosophilid flies has found a high level of integration among wing measurements (Klingenberg and Zaklan 2000). Assuming a similarly high level of integration in wing morphological traits in *T. conura*, including wing shape would allow us to compare sets of traits that differ in the degree of integration (Bolstad et al. 2014). Wings are also potentially under sexual selection in Tephritid flies, as they are used in male displays (Sivinski and Pereira 2005). To quantify wing shape in *T. conura* we used a Celestron 44308 USB microscope to take magnified images. We took one lateral image of the fly body after removal of the wings and one dorsal image of the right wing on a transparent background to allow better visibility of the wing veins. We measured body length, ovipositor length (Fig. 1D), wing length, wing width and wing area digitally from these images. Those variables were measured in units of pixels, which were then converted to units of mm using a scale that was photographed with each image. We also placed 15 landmarks, adapted from Pieterse et al. (2017; Fig. 1E) for geometric morphometrics using TPSDig2 v2.31 and TPSUtil v1.76 (Rohlf 2015). This resulted in six wing shape traits (represented by the first six principal components of x-y coordinates) for 285 female flies and 288 male flies (see Table S1 for full population sample sizes). A Procrustes fit was applied to the landmark data using PAST3 v3.20 (Hammer et al. 2001). To produce variables to include in later analyses we took the first six principal components which explained 68% of the total phenotypic variation. The melanised area of the wing was measured through an automated script developed in MATLAB (Matlab 2017) as in Nilsson et al. (2022). All subsequent statistical analyses were performed in R version 3.6.1 (R Core Team 2019).

### Measures of evolvability

Evolutionary potential – evolvability – can be measured as a mean-scaled additive genetic variance (Houle 1992). For multivariate phenotypes, evolvability measures are typically derived from mean-scaled additive genetic variance matrices (**G**) (Hansen and Houle 2008) obtained from quantitative-genetic breeding experiments. Phenotypic variance matrices (**P**) is the sum of **G** and the environmental effects shaping trait variation in the population (Steppan et al. 2002), and is sometimes used as a proxy for **G**. Although this approach has been debated (Willis et al. 1991), there is both theoretical (Cheverud 1988) and empirical (Kohn and Atchley 1988; Roff and Mousseau 2005; Porto et al. 2009; Sodini et al. 2018) evidence that **P** can be a reasonable surrogate of **G**, particularly for morphological traits (Hadfield et al. 2007). In addition, because **P** is far easier to obtain than **G, P** can be evaluated from multiple populations in their natural habitats. This allows us to study how variational properties (e.g. genetic or phenotypic variances and covariances) are related to ecology and evolve during evolutionary divergence (Berner et al. 2010; Grabowski et al. 2011; Hansen and Voje 2011; Haber 2016; Tsuboi et al. 2018). In the following analyses of P-matrices we will refer to patterns of variation as ‘evolvability’, but note that this rests on the assumption of strongly correlated environmental and genetic variation (Hansen et al. 2011).

### Estimating P-matrices

We estimated P-matrices by fitting multivariate mixed models with the `MCMCglmm` R package (Hadfield 2010) and subsequently postprocessed the posterior distributions with tools from the `evolvability` R package (Bolstad et al. 2014). For each model, we sampled the posterior distributions for 1 million MCMC iterations, with a burn-in of 500000 and a thinning interval of 500. We assumed uninformative priors for the fixed effects, and the recommended weakly informative priors for the random effect (Hadfield 2010).

All analyses were performed separately for females and males to enable us to include the functionally important ovipositor for females. To test if there are differences in evolvability between the host races, we started by estimating mean P-matrices for each host race in models including population as a random factor. To address if interpopulation variance differed between host races we fit similar models per population, but without a random factor. To disentangle if phenotypic variance differed between size, melanisation and shape traits, we repeated the analysis for three different data sets, one with all traits, one with size and melanisation traits and finally one with only wing shape traits.

To assess the distribution of variation within and among populations, we fitted separate linear mixed-effect models to log-transformed trait values, with population as a random effect. We then computed the among-population variance component as the variance among populations divided by the sum of the among-population and residual (within-population) variance.

### Comparisons of evolvability and P-matrices

To test for differences in mean evolvability between host races we first derived posterior means and credible intervals of evolvability from each estimated P-matrix, and then calculated posterior support for host plant differences in evolvability as the proportion of randomly paired posterior estimates for which evolvability was greater for the more evolvable host race than for the less evolvable host race.

To investigate if phenotypic variance differed between flies that coexist with the other host race, likely subject to some introgression (Ortega et al. *[in prep]*), and allopatric flies we performed similar comparisons of posterior means.

To assess similarity between the P-matrices estimated for the ancestral and derived host races, we correlated the expected responses to a set of random hypothetical selection gradients for the two P-matrices (Hansen & Houle 2008). This approach compares the P-matrices in terms of the parameters they are used to derive within the present theoretical framework (i.e. response to selection). We generated 1000 random selection gradients drawn from the unit sphere, and used the evolvabilityBeta function of the `evolvability` R package (Bolstad et al. 2014) to compute the evolvability along each selection gradient for each matrix.

### The effect of ancestral evolvability on divergence

Our sampling of both the ancestral and derived host race also allowed us to ask whether the derived host race has diverged in a direction of comparatively high evolvability, as expected if genetic constraints play a role in host race divergence. We asked this question in two different ways. First, we estimated a variance-covariance matrix among log-transformed population means (divergence matrix, **D**), and assessed the relationship between **D** and **P estimated for** the ancestral host race. Second, we considered the divergence of each population of the derived host race from the mean phenotype of the ancestral host race. We computed divergence vectors 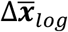 from a focal population to the mean of the ancestral host race as 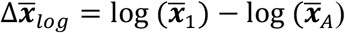, where 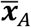 is the vector of mean phenotypes for the ancestral host race. We then computed the evolvability along 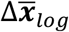 as 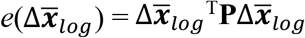, and compared this to the mean and maximum evolvability of the focal P-matrix. Divergence in a direction of greater-than-average evolvability would be consistent with some influence of ancestral variance on divergence.

### The effect of the ovipositor on evolvability and divergence

Patterns of trait divergence allowed us to formulate hypotheses about past or current patterns of selection on different traits. The ovipositor is shorter in the derived host race (Diegisser et al. 2007; Nilsson et al. 2022), and is thus likely either to be or to have been under directional selection to match the bud size of the derived host plant. The directional selection on ovipositor length in the derived host race may have depleted the variation in ovipositor length, and of the available combinations of ovipositor length and other traits within the fly populations. By comparing the autonomy, i.e. the freedom of a trait to evolve independently of other traits (Hansen and Houle 2008), of ovipositor length between the ancestral host race and the derived host race, we examined if there was evidence for a decreased autonomy in the derived host race. If the derived host race has a lower autonomy of ovipositor length than the ancestral host race, it may be an effect of reduced available variation. We further assessed if the autonomy of the ovipositor differed from the autonomies of other traits by separating overall evolvability, evolvability conditioned on all but a focal trait (hereafter referred to as overall conditional evolvability), and evolvability conditioned on ovipositor length. Overall evolvability and conditional evolvability was calculated as described elsewhere (Hansen and Houle 2008; Bolstad et al. 2014), while evolvability conditioned on ovipositor length was extracted from a separate P-matrix where we conditioned the entire matrix on ovipositor length by using the evolvabilityBeta function of the R-package `evolvability` (Bolstad et al. 2014). We also compared the autonomy of all individual traits conditioned on ovipositor length to the autonomy of the same traits conditioned to any non-ovipositor trait.

To assess whether the ovipositor plays a particular role in driving patterns of divergence, we reran the divergence-vector analyses described above using three different measures of evolvability, namely (1) raw evolvability, (2) conditional evolvability, and (3) evolvability conditioned only on ovipositor length. A clearer relationship for the latter would indicate that the ovipositor plays a specific role in driving population divergence.

## Results

Differences among populations explained an average of 14.2% of trait variance in the ancestral host race, and an average of 33.2% in the derived host race. The among-population variance component ranged across traits from 2.6% to 21.9% in the ancestral host race, and from 0.06% to 72.1% in the derived host race (Fig. 2). Some wing shape traits had too little variance to estimate variance among populations, specifically principal component 3 and 6 in the ancestral host race, and both within- and among-population variance in principal component 6 in the derived host race.

**Figure 2.**
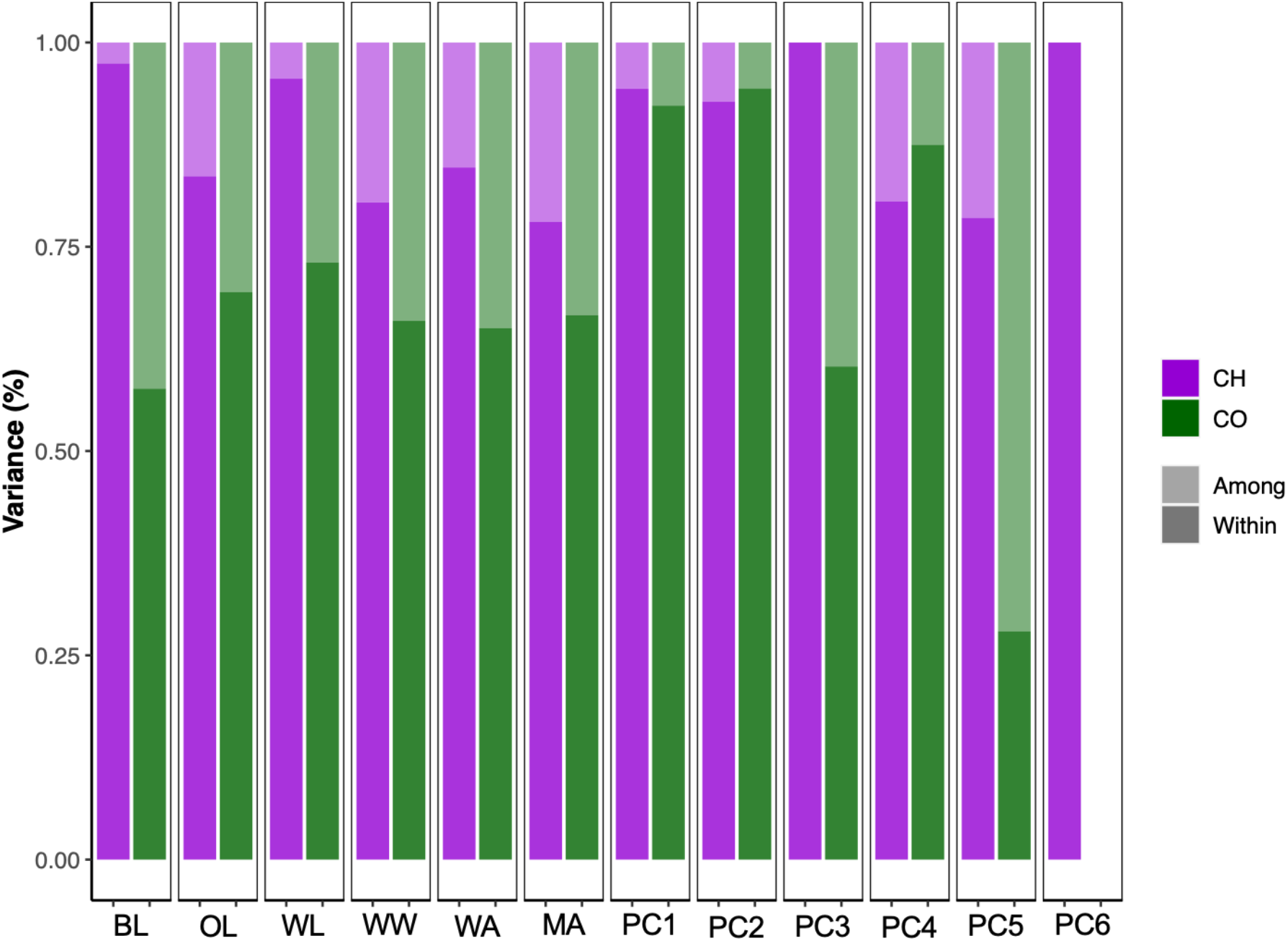
Proportional trait variance among and within populations for each host race given as percent. Note that several of the wing shape traits were insufficiently variable to disentangle among and within population variation, specifically among population variation in PC 3 and PC 6 of the ancestral host race as well as any variation in PC6 of the novel host race. Those traits are represented as zeroes here. Trait abbreviations are MA: melanised area, OL: ovipositor length, BL: body length, WW: wing width, WL: wing length and PC1-6 represents the wing shape principal components.

### Comparisons of evolvability and P-matrices

The derived host race exhibited slightly reduced evolvability compared to the ancestral host race (Fig. 3A-B). Females of the ancestral host race had a slightly broader distribution along the axis of largest variation compared to the derived host race, but the distribution along the second axis of variation was very similar for females of the two host races (Fig. 3A). In comparison, male variance differed very little between host races (Fig. 3B).

**Figure 3.**
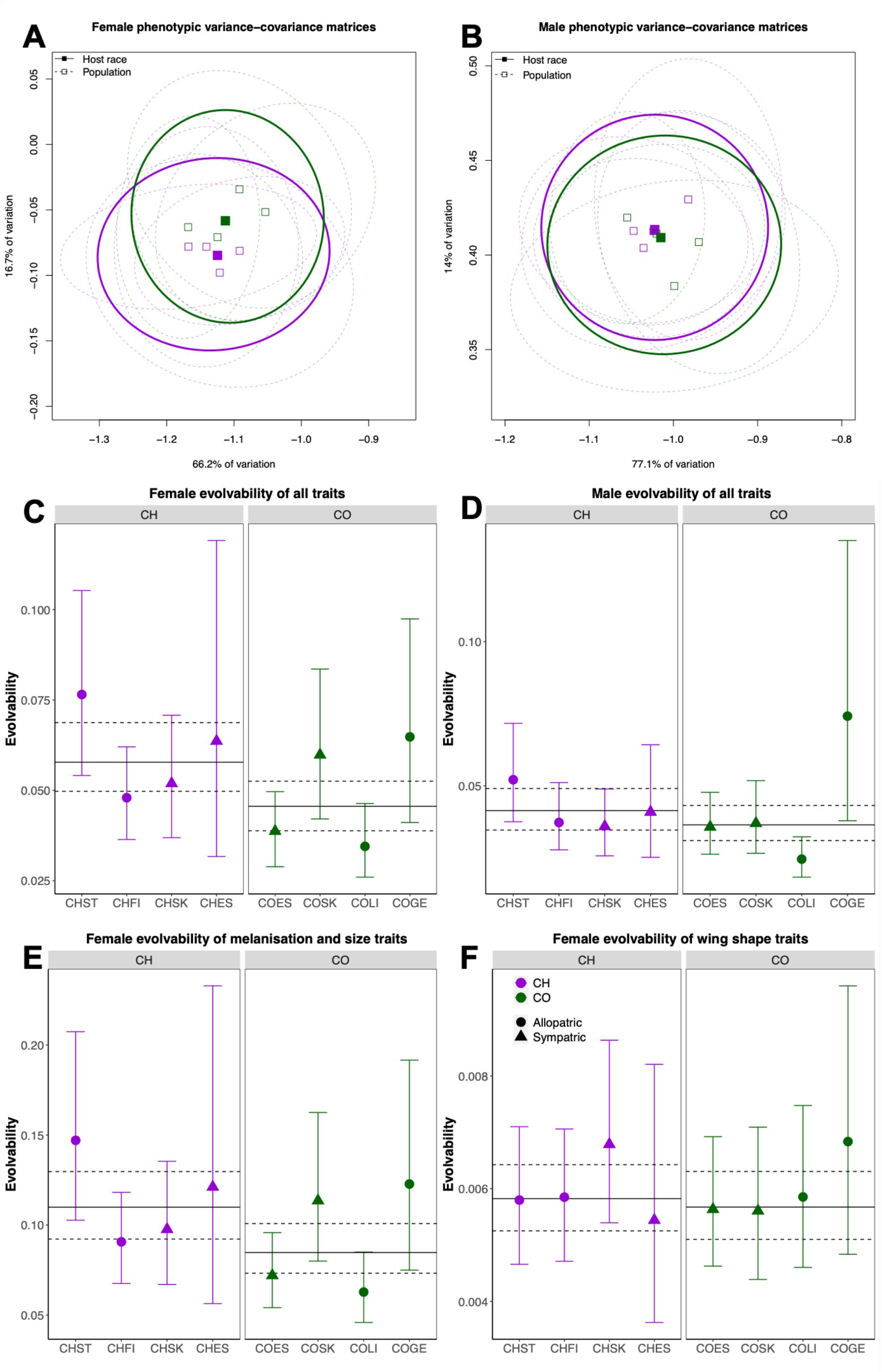
Comparisons of evolvability among populations and host races. **A)** and **B)** Two-dimensional representation of phenotypic variation based on principal components analysis on females (**A**) and males (**B**). Purple represents flies belonging to the *C. heterophyllum* host race and green represents flies belonging to the *C. oleraceum* host race (see legend Fig. 3E). Solid squares represent host race units of evolvability, and open squares represent population units of evolvability. Solid ellipses show 95% confidence interval of the overall host race measurements whereas the dashed ellipses show 95% confidence interval of the population measurements. **C** and **D)** Population and host race comparison when including all traits in females (**C**) and males (**D**). Colors are host race specific, and triangles denote sympatric populations and circles allopatric populations. Dashed lines represent maximum and minimum evolvability and solid lines represent mean evolvability as estimated by MCMC models. **E)** Population and host race comparison when including only size traits and melanisation in females. **F)** Population and host race comparison when including only female wing shape traits.

Wing shape traits were two orders of magnitude less variable than size and melanisation traits. We found small but detectable differences in evolvability between females of the two host races. Including all traits, the evolvability of females of the derived host race was 20.7% lower than that of females of the ancestral host race (posterior mean difference with SE: 1.23×10^−4^ ± 1.99×10^−6^; Fig. 3). In comparisons including size and melanisation traits only, this difference increased to 21.9% (2.48×10^−4^ ± 3.91×10^−6^). The evolvability of wing shape traits was, however, not detectably different between the host races as the decrease in evolvability in the derived host race is only 0.024% (Fig 3; Table S2).

The male p-matrices were more similar between the host races than were the female P-matrices (Fig. 3B). In turn, we failed to detect a difference in evolvability between males of the two host races for either of the trait subsets, although the evolvability was 11% lower in the derived host race compared to the ancestral host race in a comparison including both size, melanisation and wing shape traits (Fig. 3D; Table S2).

We found no support for the idea that gene flow increases evolvability, as indicated by no detectable differences in mean evolvability between allopatric and sympatric flies of either sex (Fig. 3C, 3D). Contrary to our prediction we found a 10.3% increase in evolvability in allopatric compared to sympatric populations in females when comparing all traits (Fig. 3C; Table S2), and a similar increase when including only size and melanisation traits (Fig. 3E; Table S2) whereas wing shape traits had very similar levels of evolvability in allopatric and sympatric populations (Fig. 3F; Table S2). The patterns in males mirrored those found in females (Fig. 3D; Table S2).

The ancestral and derived P-matrices were very similar, as indicated by a strong correlation between predicted selection responses to a set of random selection gradients (R^2^ = 0.97; Fig. 4A), but the relationship differs from a one-to-one slope (β = 1.42 ± 0.009). This reflects that the ancestral **P** possessed more variance among leading eigenvectors (Fig. 3A).

**Figure 4.**
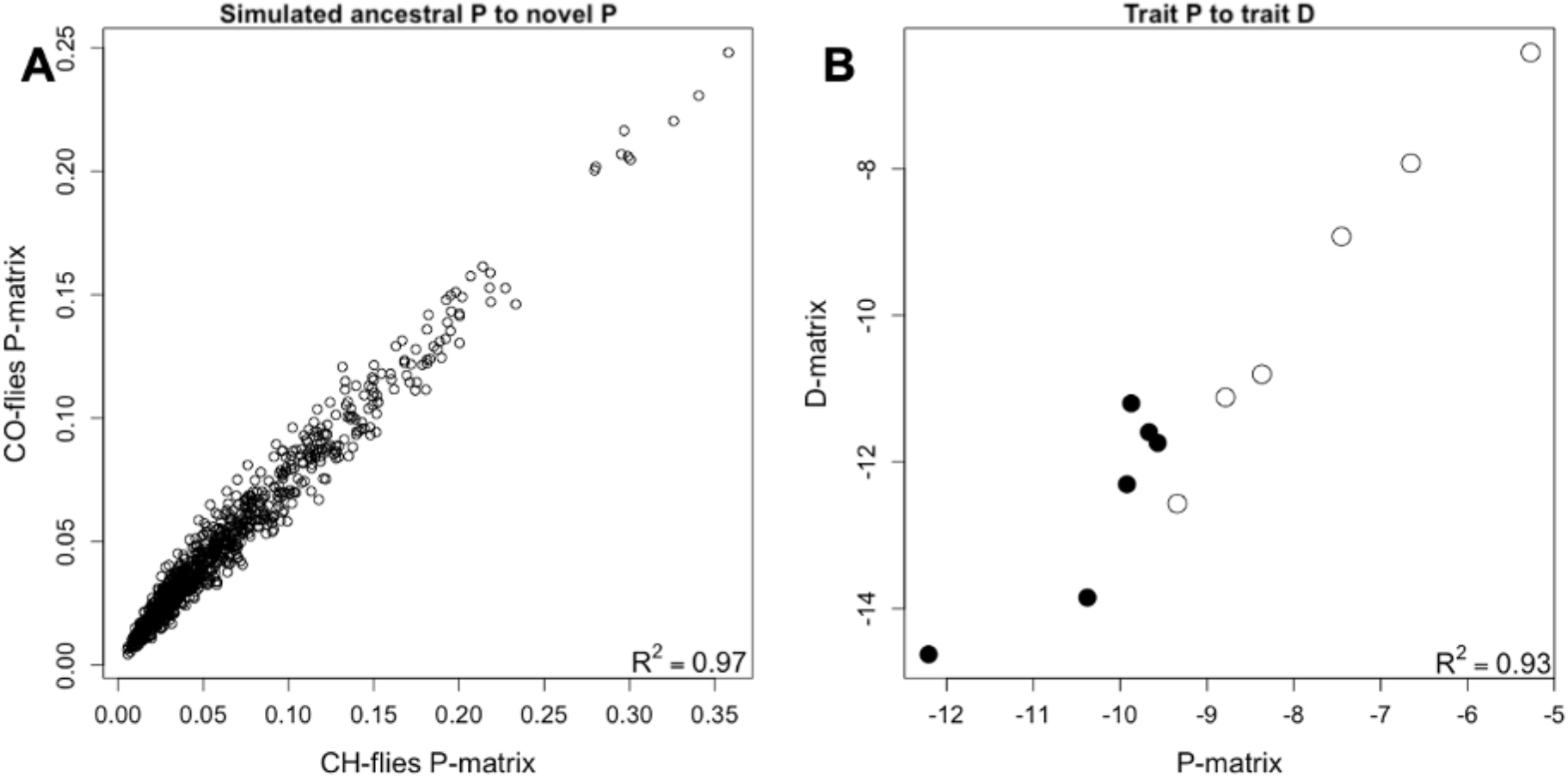
**A)** Scatterplot of variation along 1000 random selection gradients for the derived and ancestral host race. **B)** Relationship between within-population variation (P-matrix) and evolutionary divergence (D-matrix) for all traits. Open circles represent size and melanisation traits whereas closed circles represent wing shape traits.

### The effect of ancestral evolvability on divergence

To test if populations have diverged more in direction of greater ancestral evolvability, and thus how well **P** predicts divergence, we regressed evolutionary divergence between populations on variation as given by the diagonal of **P**. Size and melanisation traits line up almost perfectly (β = 1.26 ± 0.41, R^2^ = 0.97), while the relationship was somewhat less tight for wing-shape traits (β = 2.08 ± 0.67, R^2^ = 0.54; Fig. 4B).

We also asked if populations had diverged in directions of greater-than-average evolvability. This was the case for all populations (mean female population difference from overall mean with SE: 16.7% ± 0.8% for the ancestral host race and 28.2% ± 1.1% for the novel host race), and we also detected a tendency for populations diverging in directions of greater evolvability to have diverged slightly more (Fig. 5).

**Figure 5.**
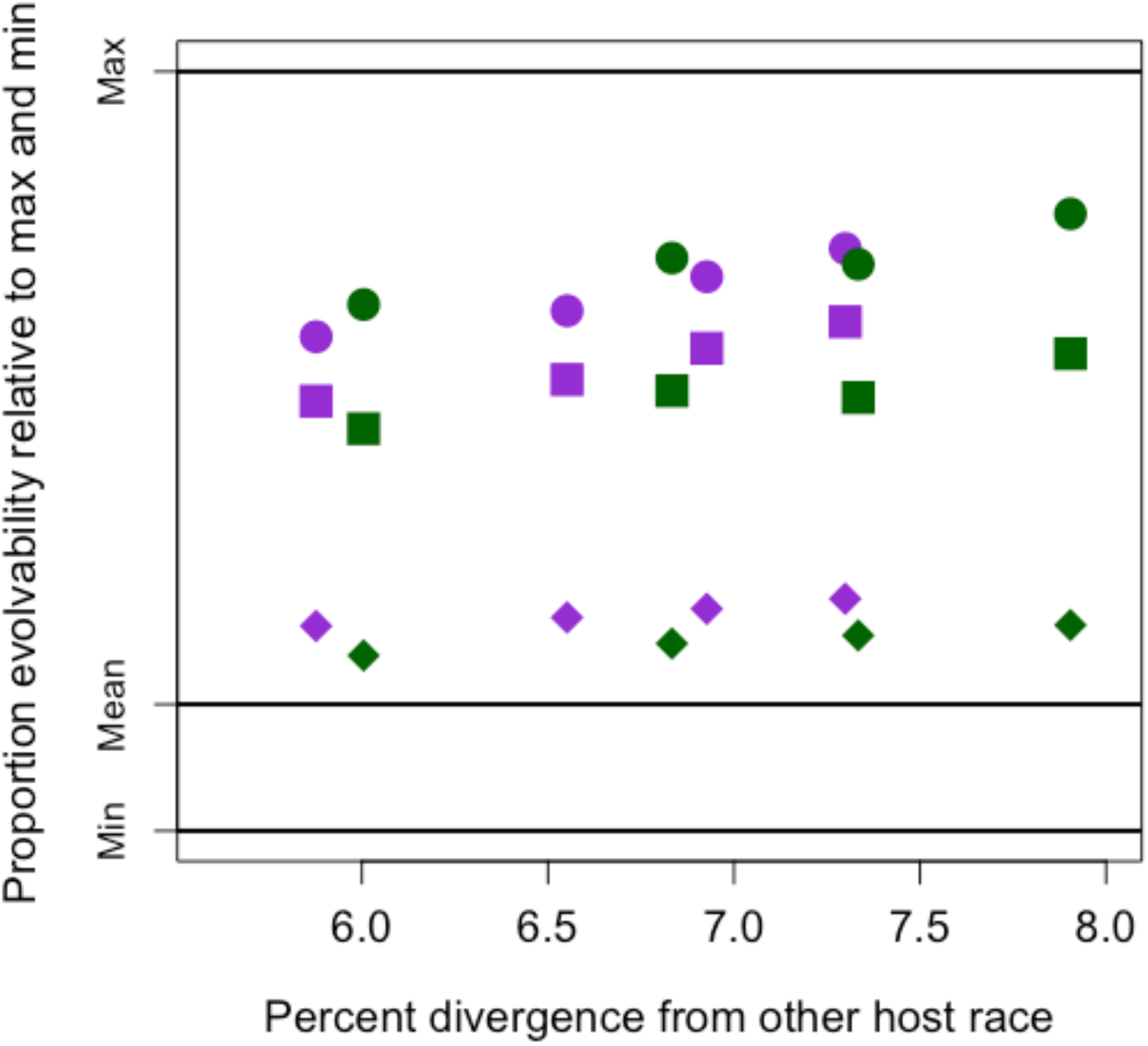
Proportional female evolvability when conditioned on ovipositor length for each population and estimated divergence from alternative host race. Full black lines represent maximum, mean and minimum evolvability of all fly females. The divergence for each population is estimated relative to the mean of all populations of the other host race. Purple points represent the ancestral host race while green points represent the novel host race. Circles represent mean evolvability, squares represent evolvability conditioned on ovipositor length and diamond shapes represent evolvability conditioned on all traits except for ovipositor length. Populations from left to right are CHES, COLI, CHFI, COES, CHST, CHSK, COGE and COSK.

### The effect of ovipositor on evolvability

Evolvability moderately reduced when conditioned only on ovipositor length, a trait we expect to have been or be under directional selection in the derived host race (Diegisser et al. 2007; Fig. 5). When comparing autonomy of ovipositor when conditioned on all traits, we found it to constrain evolution less than any other given trait (Fig. 6). Ovipositor length is thus less integrated with other traits investigated, compared to how integrated the rest of traits are with each other. Consistent with this, we found only moderate difference between overall autonomy conditioned on ovipositor and overall autonomy when excluding the ovipositor. In the ancestral females, the autonomy of a random trait conditioned on ovipositor were 74.6% of the mean autonomy, whereas it was 68.8% for the derived host race (Fig. 5).

**Figure 6.**
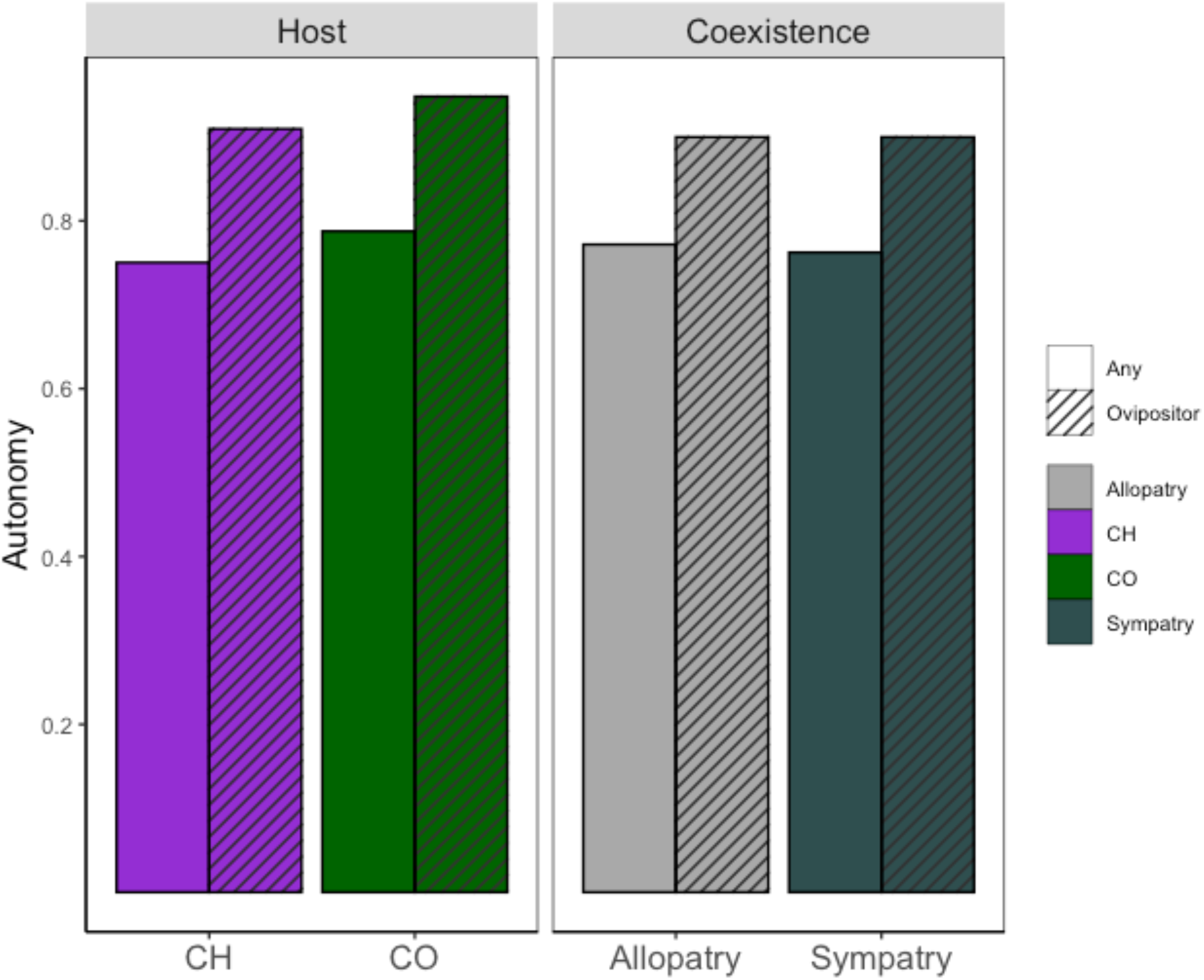
Evolvability when conditioned on ovipositor length for each of the two host races and in allopatry and sympatry. Autonomy in all traits when conditioned on any given trait or ovipositor length. Any trait represents the mean autonomy of body size, wing length, wing width, wing area and melanisation area.

## Discussion

We addressed to which extent a recent host shift, exerting directional selection on the ovipositor, had altered P-matrices in *T. conura*. We found reduced evolvability of females of the derived host race compared to the ancestral host race. This result is consistent with findings of reduced overall variation in mice that had been under artificial selection (Penna et al. 2017). One explanation for such a reduction is past or current directional selection acting on several traits following the host shift (Diegisser et al. 2006a; Diegisser et al. 2006b, 2007; Diegisser et al. 2008; Nilsson et al. 2022). Interestingly, the reduction in evolvability in the derived host race is less pronounced in males. This difference could suggest historically stronger selection on females, but this seems unlikely because there is no indication that evolvabilities of female traits are lower than those of males (Fig. 3). An alternative, and a more plausible explanation, is that the reduction reflects the inclusion of the ovipositor in the analyses of female evolvability, suggesting that ovipositor contributes to the host race differences in evolvability. The difference of female host race evolvability when excluding ovipositor length to the full set of traits is an 8.3% reduction in the ancestral host race and an 12.9% reduction in the novel host race (Table S3).

There is compelling evidence that the ovipositor is under strong selection. The length of the ovipositor is functionally important and a mismatch between ovipositor length and bud size results in reduced female reproductive success (Romstock-Volkl 1997). Therefore, the difference in ovipositor length between the host races (Diegisser et al. 2007; Nilsson et al. 2022) likely reflects adaptation to the derived host race. Historical directional selection may have caused the observed reduction in standing variation in females of *T. conura*, yet it is unclear whether there is current directional selection on the length of the ovipositor. If the population mean phenotype is already close to the new optimum associated with the new host plant, the ovipositor may currently be under stabilizing selection. Under this scenario, the reduction in variation may result from a combination of the influence of past directional and current stabilizing selection on the ovipositor, and possibly correlated responses to stabilizing selection on other morphological traits.

The influence on the ovipositor of indirect response to selection would be reduced, however, if the ovipositor is modularly independent from variation in other characters (Wagner et al. 2007; Armbruster et al. 2014). Using the concepts of conditional evolvability and autonomy, we demonstrated that this is at least partly true, because the covariance between ovipositor length and other traits reduced available variation less than did covariance among other traits (Fig. 6). This may suggest that the ovipositor constitutes a quasi-independent module separate from the other study traits. The formation of this module is not a result of the novel selective regime associated with the host shift, however, as patterns of phenotypic covariation are similar in both host races. Therefore, the existing genetic architecture may have facilitated the rapid host shift observed in *T. conura* (Diegisser et al. 2006b) by allowing divergence in ovipositor length without disrupting other traits.

One possible explanation for the reduced variation in the derived host race is that a stringent bottleneck event occurred at the host transition phase, as suggested by the lower variation in mitochondrial haplotypes in the derived host race (Diegisser et al. 2006b). This would substantially reduce the variation available within the ancestral gene pool (Nei et al. 1975), and potentially evolvability. Moreover, the host races overlap in phenology by 16% (Romstock-Volkl 1997), and larval survival on the alternative host plant is 10% (Diegisser et al. 2008). Thus only 1.6% of individuals are expected to survive on the alternative host plant, given a uniform phenotypic distribution and random mating among host races. The host shift has reduced the size of **P** in the derived host race, especially for females, while our analyses suggest limited changes in the shape of **P** (Fig. 3 A-B). Our findings add to the evidence that **P** (or **G**) may change following divergence (Eroukhmanoff and Svensson 2011; Björklund et al. 2013), although the changes in **P** we find are moderate and sex-dependent. Our findings may be inherent to the traits we decided to measure, or an effect of using **P** as a proxy for **G**. Our use of **P** as a proxy for **G** is justified based on previous case studies (Kohn and Atchley 1988; Roff and Mousseau 2005; Porto et al. 2009), empirical assessment (Sodini et al. 2018) and theoretical work (Cheverud 1988). The *Tephritidae* family is relatively closely related to *Drosophilidae*, a family where **P** has been found to approximate **G** (but see McGuigan and Blows 2007).

Our chosen size and melanisation traits had orders of magnitude higher evolvability than wing shape traits. Low variance in the shape compared to size and melanisation traits in *T. conura* is consistent with other studies reporting shape to be less evolvable than size (Hunt 2007; Houle et al. 2017). At a glance, this result contrasts with our previous finding that wing shapes differ between *T. conura* host races (Nilsson et al. 2022). Evolvability is a measure of expected response to selection in percent of the trait mean following an episode of unit strength selection (Houle 1992). Despite a comparatively low evolvability of in wing shape, that is approximately 0.006% of centroid size (Fig. 3F), the half-time, i.e. the number of generations needed to double or halve the trait measurement, could be rather quick in an evolutionary time scale. Given an assumed standard heritability of 0.27 (estimate of shape traits from Hansen and Pelabon (2021)) the half-time would be roughly 43 thousand generations under persistent directional selection. Given the divergence time between the derived and ancestral host specialists estimated to have coincided with the glacial retraction following the most recent ice age (Diegisser et al. 2006a) and the univoltine nature of *T. conura*, our estimates of evolvability could result in appreciable variation in wing shape. Therefore, the lower evolvability of wing shape compared to size traits should not be taken as evidence of low evolvability at evolutionary time scales.

One reason for low evolvability in wing shape of *T. conura* is that these traits are tightly integrated and related to overall wing size. The Procrustes fit we applied to the wings standardize the size and alignment of all images was aimed to compare shape while correcting for size. If we instead perform a Procrustes fit without scaling wing shape to centroid size, 97% of total variance is explained by the first principal component, representing wing size. Almost all of the variation in wing shapes are strongly correlated to size variation (Fig. S3), which we removed by correcting to centroid size. This suggests that the variation in wing morphology of *T. conura* is largely a matter of scaling up and down of the exact same wing shape. Such isometric scaling is in contrast with wing shape in another Dipteran family *Drosophilidae*, which shows considerable shape variation that are unrelated to size (Bolstad et al. 2015). In the future, it would be interesting to investigate if wing shape is isometrically related to size in other *Tephritidae* flies.

An additional factor that could affect evolvability is gene flow among the host races, as gene flow could increase the available genetic variation, and thereby increase evolvability in sympatry (Blows and Higgie 2003; Dochtermann and Matocq 2016; Gompert et al. 2017). Contrary to this prediction, allopatric and sympatric populations had similar levels of evolvability. Thus, there is no indication that gene flow affects evolvability in *T. conura*, but we would need formal tests of introgression in sympatric populations to ascertain the validity of this conclusion. One interpretation of this result is that novel genetic variants introduced by gene flow does not necessarily have phenotypic consequences. This may be the case particularly because the size and shape traits in our study most likely have highly polygenetic genetic architecture (Noble et al. 2017). Alternatively, the lack of effects on the evolvability of *T. conura* could be explained by the age of secondary sympatry, as evolvability is predicted to increase following early gene flow due to linkage disequilibrium, but may be reduced to normal levels after a time of recombination (Tufto 2000). Alternatively, genetic drift may play a role in the lack of differences between sympatric and allopatric populations, as several of the sympatric populations were sampled from the edges of the distributions of the *T. conura* host races. There, effective population sizes may be smaller than in range center populations. Genetic drift affects the phenotype (**P)** but in stochastic ways (Roff and Mousseau 2005), and a comparison of **P** and neutral sites would need to be performed to assess the effects of drift on *T. conura* evolvability.

There is a possibility that phenotypic plasticity could contribute to our findings. Although the flies were hatched in a standardized environment, host plant and sampling specific effects could nevertheless be expected. There are, however, at least two sympatric and two allopatric populations of each host race sampled, implying that each type of population experienced at least two different environments. Furthermore, the strong correlation between **P** and divergence between the host races (Fig. 4B and 5) does, however, suggest that **P** is an encouragingly reasonable approximation of **G** in *T. conura*.

## Conclusions

We find evidence for reduced current evolvability in response to past or current directional selection resulting from the colonization of a new niche in females, but not males, of *T. conura*. Potentially, the differences between sexes could suggest that selection on the ecologically important ovipositor is responsible for the observed reduction in evolvability. Our study adds to growing evidence that evolvability is predictive of divergence between populations (Bolstad et al. 2014; Houle et al. 2017; McGlothlin et al. 2018; Opedal et al. 2023), and illustrates that evolvability is a dynamic entity that evolves when populations are exposed to novel environments.

## Acknowledgements

We thank Mikkel Brydegaard for his assistance in programming the melanisation trait measurements, Jesús Ortega, Jodie Lilley, Emma Kärrnäs, and Mathilde Schnuriger for help in the field and in the lab. This study was funded by a Wenner-Gren Fellowship, a Crafoord grant and a Swedish Research Council grant to A. R..

