## Supplemental Material for "Colonization of a novel host plant reduces phenotypic variation"

Table S1: Locations and sample size of *T. conura* populations used.

| Population | Country | Host Plant | Co-existence | Transect | Female n | Male n |
|---|---|---|---|---|---|---|
| CHES | Estonia | *C. heterophyllum* | Sympatric | East | 16 | 23 |
| CHFI | Finland | *C. heterophyllum* | Allopatric | East | 47 | 47 |
| CHST | Sweden | *C. heterophyllum* | Allopatric | West | 42 | 46 |
| CHSK | Sweden | *C. heterophyllum* | Sympatric | West | 40 | 32 |
| COES | Estonia | *C. oleraceum* | Sympatric | East | 47 | 50 |
| COLI | Lithuania | *C. oleraceum* | Allopatric | East | 36 | 39 |
| COGE | Germany | *C. oleraceum* | Allopatric | West | 22 | 17 |
| COSK | Sweden | *C. oleraceum* | Sympatric | West | 35 | 34 |

Table S2: Posterior support from MCMC models comparing mean evolvability between groups. Both comparisons between host races, in different modes of coexistence are shown, for both sexes.

| Test | Sex | Traits | Percent difference | Mean difference | 2.5% percentile | 97.5% percentile | p |
|---|---|---|---|---|---|---|---|
| Hostrace | Female | All traits | 20.7 | $1.23 \times 10^{-4}$ | $4.12 \times 10^{-6}$ | $2.53 \times 10^{-4}$ | 0.02 |
| Hostrace | Female | No shape | 21.9 | $2.48 \times 10^{-4}$ | $1.73 \times 10^{-5}$ | $4.84 \times 10^{-4}$ | 0.01 |
| Hostrace | Female | Only shape | 0.02 | $1.55 \times 10^{-6}$ | $-6.79 \times 10^{-6}$ | $1.05 \times 10^{-5}$ | 0.37 |
| Co-existence | Female | All traits | 10.3 | $-4.70 \times 10^{-5}$ | $-1.70 \times 10^{-4}$ | $7.04 \times 10^{-5}$ | 0.23 |
| Co-existence | Female | No shape | 10.4 | $-8.96 \times 10^{-5}$ | $-3.18 \times 10^{-4}$ | $1.40 \times 10^{-4}$ | 0.23 |
| Co-existence | Female | Only shape | 0.75 | $-2.58 \times 10^{-7}$ | $-8.46 \times 10^{-6}$ | $8.63 \times 10^{-6}$ | 0.48 |
| Hostrace | Male | All traits | 11 | $4.86 \times 10^{-5}$ | $-4.58 \times 10^{-5}$ | $1.47 \times 10^{-4}$ | 0.15 |
| Hostrace | Male | No shape | 10.3 | $9.18 \times 10^{-5}$ | $-9.90 \times 10^{-5}$ | $3.06 \times 10^{-4}$ | 0.2 |
| Hostrace | Male | Only shape | 18.2 | $1.29 \times 10^{-5}$ | $2.56 \times 10^{-6}$ | $2.25 \times 10^{-5}$ | 0.01 |
| Co-existence | Male | All traits | 15.8 | $-5.45 \times 10^{-5}$ | $-1.43 \times 10^{-4}$ | $3.83 \times 10^{-5}$ | 0.11 |
| Co-existence | Male | No shape | 16.6 | $1.14 \times 10^{-4}$ | $-3.15 \times 10^{-4}$ | $6.25 \times 10^{-5}$ | 0.13 |
| Co-existence | Male | Only shape | 11 | $-2.79 \times 10^{-6}$ | $-1.29 \times 10^{-5}$ | $7.64 \times 10^{-6}$ | 0.27 |

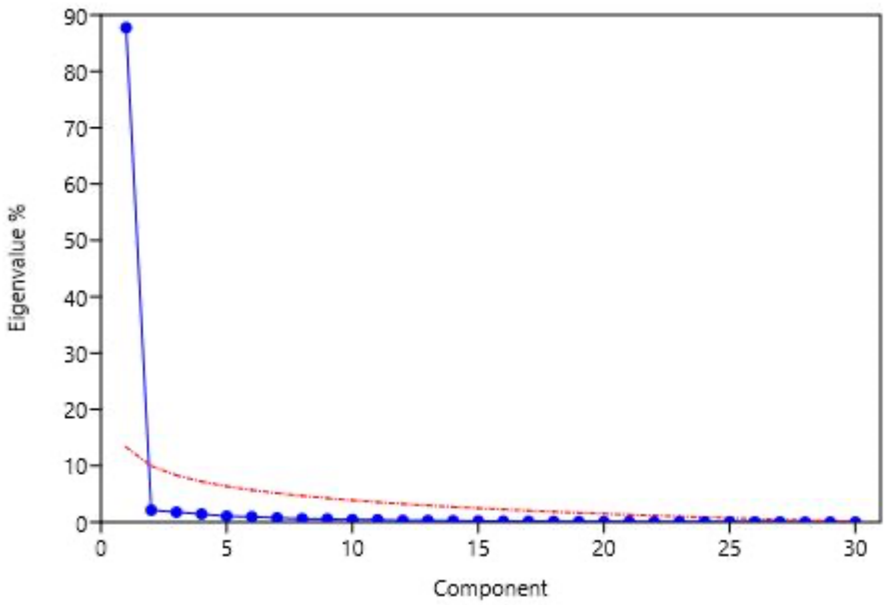


Figure S1: Scree plot describing eigenvalues of wing shape traits without correcting for size.

A large majority of variation is captured by the first principal component, i.e. size.

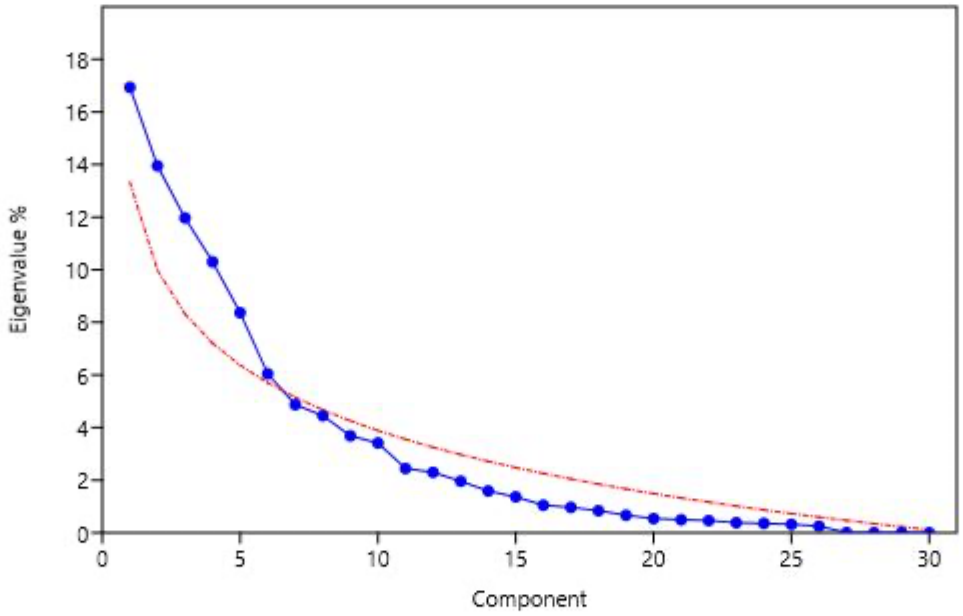


Figure S2: Scree plot describing eigenvalues of wing shape traits after correcting for size. The six first principal components shown here above the red ‘broken stick’-line are the principal components we used as wing shape traits throughout the analysis.

Table S3: Posterior support from MCMC models comparing mean evolvability within host races between original datasets compared to datasets with ovipositor length removed. Only females included.

| Test | Host race | Percent difference | Mean difference | 2.5% percentile | 97.5% percentile | p |
|---|---|---|---|---|---|---|
| With and without ovipositor | Ancestral | 8.32 | $5.19 \times 10^{-5}$ | $-7.76 \times 10^{-5}$ | $1.81 \times 10^{-4}$ | 0.23 |
| With and without ovipositor | Novel | 12.9 | $6.14 \times 10^{-5}$ | $-3.51 \times 10^{-5}$ | $1.63 \times 10^{-4}$ | 0.11 |
